# FRAGMENT MORPHOMETRY ANALYSIS AND SAME-COLOR-CHANNEL SEPARATION ENABLE OBJECTIVE QUANTIFICATION ACROSS BBB MODELS

**DOI:** 10.64898/2026.06.26.734824

**Authors:** Benjamin D. Peck, Nicholas R. O’Hare, Craig F. Ferris, Rebecca L. Pinals, Eno E. Ebong

## Abstract

Quantifying blood-brain barrier (BBB) integrity from fluorescence microscopy remains limited by subjective scoring and categorical classification methods that lack reproducibility. For objective and consistent BBB phenotyping, we present two semi-automated image-analysis pipelines that replace manual scoring with quantitative, continuous-variable measurements.

Our *in vitro* pipeline, implemented in Python, quantifies the connectivity of tight junction structures by measuring discrete ZO-1 fragment objects within manually traced junction regions. It outputs continuous metrics including average fragment area, total junctional area, and a junctional fragmentation ratio that captures degree of ZO–1 continuity versus discontinuity. In human brain microvascular endothelial cells subjected to glycocalyx component knockdown, the pipeline detected significantly reduced fragment area (37% decrease for both CD44 and syndecan-1 (SDC1) knockdown, p = 0.0148 and 0.0084) and junctional fragmentation ratio (p = 0.0061 and 0.0137).

Our *in vivo* pipeline integrates ilastik-based pixel classification with FIJI macro automation to quantify vascular marker colocalization and to separate vessel signal from microglial contamination within a single fluorescence channel, eliminating the need for dedicated counterstains. Applied across four mouse cohorts [young, aged, Alzheimer’s, traumatic brain injury (TBI)] and three brain regions [prefrontal cortex (PFC), hippocampus, midbrain], the pipeline revealed concurrent ZO-1 loss and ICAM-1 elevation in the PFC and hippocampus of aged and Alzheimer’s mice, with Alzheimer’s-specific doubling of eNOS occurring in the PFC (p = 0.0013). TBI mice showed persistent ZO-1 loss with transient ICAM-1 and eNOS changes.

Both deterministic pipelines are available on GitHub and designed for adoption beyond the specific markers and systems analyzed here.

## 1. INTRODUCTION

Blood-brain barrier (BBB) dysfunction is observed across neurological conditions, including Alzheimer’s disease (AD), traumatic brain injury (TBI), and normal aging [1–3]. In the case of TBI, acute mechanical injury directly damages the vasculature [1], and in normal aging, gradual vascular deterioration compromises barrier integrity over time [2, 4]. Whether or not vascular changes precede other pathological features in each condition, detecting and quantifying BBB alterations is important for understanding when and how it fails. BBB integrity encompasses both structural features, such as tight junction protein expression and organization, and functional properties, such as transendothelial permeability. However, existing methods for quantifying these BBB-associated markers rely on subjective scoring or categorical classification, and no published, standardized pipeline exists for the *in vivo* setting. This work addresses the gap by establishing a highly sensitive, reproducible image-analysis framework for quantifying BBB-associated vascular markers and resolving subtle differences between conditions.

### 1.1 ALZHEIMER’S DISEASE AND THE VASCULAR HYPOTHESIS

AD is the most common form of dementia, characterized by progressive memory loss, cognitive decline, and neuronal death [5]. The main pathological hallmarks of AD are amyloid-beta (Aβ) plaques and neurofibrillary tangles (NFTs). Aβ plaques are extracellular aggregates of the Aβ peptide, which is normally cleared from the brain but accumulates pathologically and disrupts synaptic signaling [5, 6]. NFTs are abnormal aggregates of tau, a microtubule-associated protein that normally stabilizes the internal structure and intracellular transport network of neurons. NFTs accumulate within nerve cells and disrupt the delivery of essential molecules to synapses [7].

Aβ is produced by enzymatic cleavage of amyloid precursor protein (APP) and exists in two major isoforms, Aβ40 and Aβ42, which differ in their aggregation properties and predominant sites of accumulation. Aβ42 is the dominant species in parenchymal plaques, while Aβ40 is enriched in and around blood vessel walls in a condition known as cerebral amyloid angiopathy, though both isoforms are found in both compartments [8]. Aβ is cleared from the brain through multiple routes, including transport across the vasculature, microglial phagocytosis, and glymphatic drainage [9]. When these clearance mechanisms fail or Aβ production is excessive, both isoforms accumulate and form toxic aggregates. For decades, research and drug development have focused on clearing these protein aggregates, but even recent anti-amyloid antibodies that achieve measurable plaque clearance, such as lecanemab (which achieved 27% slowing of cognitive decline [10]) and donanemab (which achieved 35% reduction in progression risk [11]), have produced only modest clinical benefit.

The limited success of these approaches has led researchers to look beyond amyloid for earlier disease drivers. The vascular hypothesis of AD proposes that dysfunction of the brain’s blood vessels is not a late consequence of protein accumulation, but an early event that precedes and facilitates it [4, 6]. Under this framework, it is hypothesized that BBB breakdown impairs the clearance of Aβ from the brain while simultaneously permitting the influx of circulating neurotoxic factors, initiating a self-reinforcing cycle of vascular damage and protein accumulation that may ultimately drive neurodegeneration [6]. Transcriptomic profiling of brain vasculature across 101 individuals spanning the AD continuum has shown that endothelial cells exhibit some of the earliest and most pronounced transcriptional changes, detectable as early as mild cognitive impairment [12]. Detecting these changes at the protein level requires quantitative imaging tools capable of measuring vascular marker expression in tissue.

### 1.2 THE BLOOD-BRAIN BARRIER

The BBB is a multicellular vascular interface that regulates the exchange of molecules and cells between the blood and brain [2, 3]. It is formed by brain microvascular endothelial cells (ECs), which line the interior of cerebral blood vessels, together with pericytes and astrocytes that provide structural and signaling support [2, 3]. Adjacent ECs are connected by tight junction complexes, which are protein assemblies that seal the paracellular space between neighboring cells and tightly regulate molecular transport into the brain [3]. The transmembrane components of these complexes include claudins (particularly claudin-5) and occludin, which span the cell membrane and directly contact the corresponding proteins on the neighboring cell [3, 13]. The scaffolding protein zonula occludens-1 (ZO-1) anchors these transmembrane proteins to the intracellular actin cytoskeleton and is critical for maintaining tight junction integrity [14, 15]. In intact endothelium, ZO-1 appears as continuous bands at cell-cell borders in fluorescence microscopy, and under barrier compromise, ZO-1 bands break down into multiple discrete fragments [13, 16]. The degree of fragmentation is representative of the severity of junction disruption, which motivates the fragment-level morphometric analysis developed in this study. Vascular endothelial cadherin (VE-cadherin) mediates adherens junctions, a second type of cell-cell junction that mechanically links neighboring ECs and helps maintain the structural integrity of the vessel wall. Unlike ZO-1, which is also present diffusely in the cytoplasm, VE-cadherin localizes specifically to cell-cell contacts, making it a reliable reference for identifying junctional regions in microscopy [17].

On the luminal (or blood-facing) side of the ECs, the endothelial glycocalyx (GCX) forms the outermost interface between the BBB and circulating blood [18]. The GCX is a carbohydrate-rich layer composed of core proteins, including cluster of differentiation 44 (CD44) and syndecan-1 (SDC1), and their associated sugar chains [18]. The GCX mediates endothelial sensing of blood flow shear stress (mechanotransduction), regulates vascular barrier stability and permeability, and mitigates inflammation [18–20]. Because the GCX’s role in BBB regulation is incompletely defined, we in the Ebong laboratory developed an *in vitro* BBB model for mechanistic studies of glycocalyx-tight junction interactions [21]. Using this model, ongoing work is examining how cultured human brain microvascular endothelial cell (HBMEC) loss of specific GCX components, including CD44 and SDC1, affects ZO-1 expression and barrier permeability. These GCX knockdown (KD) conditions serve as the experimental basis for the *in vitro* fragmentation analysis in this study, providing a controlled system in which tight junction disruption has been confirmed by complementary permeability measurements.

### 1.3 VASCULAR MARKERS OF BBB HEALTH

This study assesses ZO-1 as a key dimension of BBB health. ZO-1 immunofluorescence is one of the most commonly used readouts for evaluating tight junction integrity in both cell culture and brain tissue [3, 13, 14, 17]. As stated previously, ZO–1 shifts from continuous junctional bands to discrete segments when barrier organization changes, providing a compact structural readout of tight junction integrity.

To capture endothelial inflammatory status alongside tight junction structure, this study quantifies ICAM–1 as a complementary marker to ZO-1. ICAM-1 is a transmembrane glycoprotein that mediates leukocyte adhesion to and transmigration across the endothelium [20]. It is expressed at low levels on healthy endothelium under non-inflammatory conditions and upregulated in response to inflammatory cytokines and oxidative stress [22, 23]. Elevated ICAM-1 at the BBB has been documented in AD, TBI, and aging [1, 23], making it a relevant indicator of endothelial activation associated with these conditions. Because ICAM-1 upregulation occurs on the endothelial surface, it can be quantified as a coverage metric on lectin-defined vessel boundaries using the colocalization pipeline described in Section 2.4.

To further assess endothelial functional state, this study also quantifies endothelial nitric oxide synthase (eNOS), which catalyzes the production of nitric oxide (NO), a regulator of vascular permeability [24] and cerebral blood flow through vasodilation [25]. At physiological levels, eNOS-derived NO helps maintain endothelial barrier integrity by stabilizing VE-cadherin at cell-cell junctions, and eNOS is the predominant NO synthase isoform controlling vascular permeability *in vivo* [26]. However, under pathological conditions such as oxidative stress, eNOS can become uncoupled, producing reactive oxygen species instead of NO [27]. This shifts the balance from NO-mediated barrier protection to oxidative barrier damage [24]. eNOS activity is also modulated in part by the GCX, which helps mediate its shear stress-dependent activation [19]. eNOS coverage on vessel boundaries therefore provides a readout of endothelial phenotype across disease conditions.

### 1.4 DISEASE MODELS AND BRAIN REGIONS

The analyses in this study span four mouse cohorts representing distinct BBB challenges: young healthy controls (3 months, C57BL/6), mild repetitive TBI (3 months, C57BL/6), aged (18 months, C57BL/6), and AD (18 months, APPswe/PSEN1dE9 on C57BL/6 background) [5]. The APPswe/PSEN1dE9 model is a widely used and well-established genetically modified model of familial AD, carrying mutations in both amyloid precursor protein and presenilin-1 [5, 8]. The PSEN1dE9 mutation shifts Aβ production toward Aβ42, so the APPswe/PSEN1dE9 model develops extensive parenchymal plaque deposition but limited cerebral amyloid angiopathy [8]. The aged and AD cohorts are age-matched at 18 months, enabling AD-specific pathology to be distinguished from normal aging.

TBI serves as a positive control for acute BBB disruption [28–30], reliably inducing tight junction protein loss including ZO-1 [31], and transient upregulation of both endothelial ICAM-1 [32] and eNOS [33, 34]. TBI cohort data are therefore assessed qualitatively to confirm that the pipeline detects expected marker changes rather than subjected to formal statistical analysis.

Three brain regions were selected for analysis based on their differing vulnerability to age- and disease-related vascular changes. The prefrontal cortex (PFC) is involved in executive function and decision-making and shows early BBB changes in both aging and AD [35]. The hippocampus is central to memory formation and is among the earliest regions affected in AD, with well-documented susceptibility to age-related vascular dysfunction [4, 36]. The midbrain, which supports motor coordination and sensory processing, is relatively spared from Aβ deposition in the APPswe/PSEN1dE9 model and was included as a less-affected reference region [37]. Comparing vascular marker expression across these three regions allows region-specific BBB changes to be separated from global ones.

### 1.5 LIMITATIONS OF CURRENT QUANTITATIVE METHODS

Despite the widespread use of ZO-1 staining for evaluating tight junction integrity, current analysis of junction morphology remains largely qualitative. Investigators typically score junctions as “continuous,” “discontinuous,” or “absent” by manual inspection [13], and classification-based tools such as the JAnaP formalize this approach by sorting junction segments into morphological categories along manually traced cell perimeters [17]. Other tools quantify junction organization without categorical scoring: TiJOR captures intensity variation along junction contours [38], IJOQ estimates continuity through grid-based intersection counting [39], and Junction Mapper measures junction area and intensity along defined cell-cell interfaces but does not resolve disconnected fragments within those interfaces [40]. Lu et al. applied connected component analysis to claudin-7 in tissue sections [41], identifying fragments as discrete objects, but reported only a global fragmentation index per image rather than measuring individual fragment area, perimeter, or shape. None of these tools produce per-fragment morphometric measurements that would allow the size distribution and geometry of junction fragments to be characterized across experimental conditions.

For *in vivo* vascular marker analysis, general-purpose image processing tools such as FIJI provide the individual operations needed to quantify marker expression (thresholding, masking, intensity measurement) [42], but no published, standardized workflow exists for applying them reproducibly across brain regions and disease cohorts, and individual laboratories typically develop custom macros that are not shared or validated externally.

The *in vivo* setting also introduces a technical challenge that does not arise in cell culture. Cerebral vasculature in tissue sections is commonly labeled using tomato lectin, which binds sugar residues on endothelial cell surfaces [43, 44]. However, activated microglia (the brain’s resident immune cells) also express surface glycoproteins bearing the same sugar residues [43, 44], and in neuroinflammatory conditions this microglial signal contaminates the vessel label and distorts all downstream measurements. Conventional intensity thresholding cannot separate the two cell types because both appear as bright objects in the same channel. A dedicated microglial counterstain such as Iba1 could resolve the ambiguity, but in multi-marker experiments the available fluorophore channels are often fully committed to the markers of interest, and when the protein of interest and the microglial marker require primary antibodies raised in the same host species, both cannot be used simultaneously because the secondary antibody would bind to both primary antibodies indiscriminately. Machine learning approaches have been applied to segment microglia and vasculature independently in dedicated fluorescence channels [45, 46], but these methods operate on channels where only one cell type is labeled. When both cell types are labeled by the same marker within a single channel, as occurs with tomato lectin in neuroinflammatory tissue, no existing pipeline separates them without a dedicated counterstain.

This study addresses these gaps with two image analysis pipelines. The first, implemented in Python, quantifies ZO-1 tight junction fragmentation *in vitro* through adaptive background correction and fragment-level morphometric analysis. The second combines ilastik-based machine learning with FIJI macro automation to quantify vascular marker expression within lectin-defined vessel boundaries *in vivo*, including a classification step that separates vessel and microglial signal within a single fluorescence channel without a dedicated microglial stain. Both pipelines are applied to the experimental systems described above and are publicly available on GitHub.

## 2. METHODS

### 2.1 *IN VITRO* IMAGE DATASET

Cell culture, GCX KD experimentation, immunofluorescence staining, and confocal imaging were conducted as part of ongoing parallel studies. Briefly, primary HBMECs (Cell Systems; passages 4-8) were exposed to 12 dyne/cm² shear stress for 24 hours using a custom millifluidic flow system [21]. Stable GCX KD was achieved using shRNA-containing lentiviral particles targeting CD44 and SDC1 (OriGene). A non-targeting scrambled shRNA construct expressing GFP served as the control, accounting for any effects of lentiviral transduction itself. Cells were stained for ZO-1 (Alexa Fluor 555) and VE-cadherin (VE-CAD, Alexa Fluor 647) and imaged on a Zeiss LSM 880 confocal microscope at 63x magnification with 1.4 numerical aperture oil-immersion objective.

In the exported composite images, ZO-1 was displayed in green and VE-cadherin in red. All subsequent references to channel colors refer to these display assignments, not the fluorophore emission wavelengths. Images were acquired as 512 x 512-pixel confocal Z-stacks and stored in Zeiss CZI format, the proprietary file format used by Zeiss confocal microscopes. Five independent flow experiments per condition, with three fields of view imaged per experiment, yielded n = 15 images per condition (n = 45 total). One CD44 KD experiment was excluded prior to analysis because the junctional fluorescence was too dim to trace reliably, leaving four experiments (12 images) for that condition. This exclusion was made prior to analysis based solely on insufficient ZO-1 and VE-CAD signal in the composite image. These images served as the input dataset for the ZO-1 fragmentation analysis pipeline described in Section 2.2.

### 2.2 DEVELOPMENT OF ZO-1 FRAGMENTATION ANALYSIS PIPELINE

The ZO-1 fragmentation analysis pipeline was implemented in Python 3.12.3 using open-source image processing libraries: bioio for reading microscopy file formats, NumPy and SciPy for numerical computation, scikit-image for image segmentation and morphometric measurement, OpenCV and Pillow for image manipulation, pandas for tabular data organization, and matplotlib for visualization. The complete source code has been made available as a Jupyter notebook (an interactive computing environment that combines code, visualizations, and documentation in a single file) on GitHub (https://github.com/peckbe/junction-fragmentation-analysis) under the MIT license.

The pipeline was organized as three Jupyter notebooks. The first loaded CZI files and exported composite images for manual tracing. The second imported the completed tracings and matched them to their corresponding CZI files. The third performed all quantitative analysis: adaptive background correction, fragment identification, morphometric measurement, and data export. Only the tracing step between the first and second notebooks required human input. All analysis in the third notebook was fully automated, deterministic, and reproducible.

#### 2.2.1 Image Loading and Maximum Intensity Projection

Confocal Z-stacks contain multiple focal planes through the cell monolayer, but the fragmentation analysis operates in two dimensions. The first notebook in the pipeline loaded each CZI file and collapsed each channel’s Z-stack into a single image via maximum intensity projection, retaining the brightest value at each pixel position across all focal planes. The resulting ZO-1 channel (**Figure 1A**) and VE-cadherin channel were combined into a composite RGB image (**Figure 1B**) in which both channels were visible simultaneously, allowing junctions to be identified by VE-cadherin localization independent of ZO-1 morphology. These composite images were exported for manual junction tracing. Pixel size was read automatically from the CZI file metadata and stored for downstream unit conversion.

**Figure 1:**
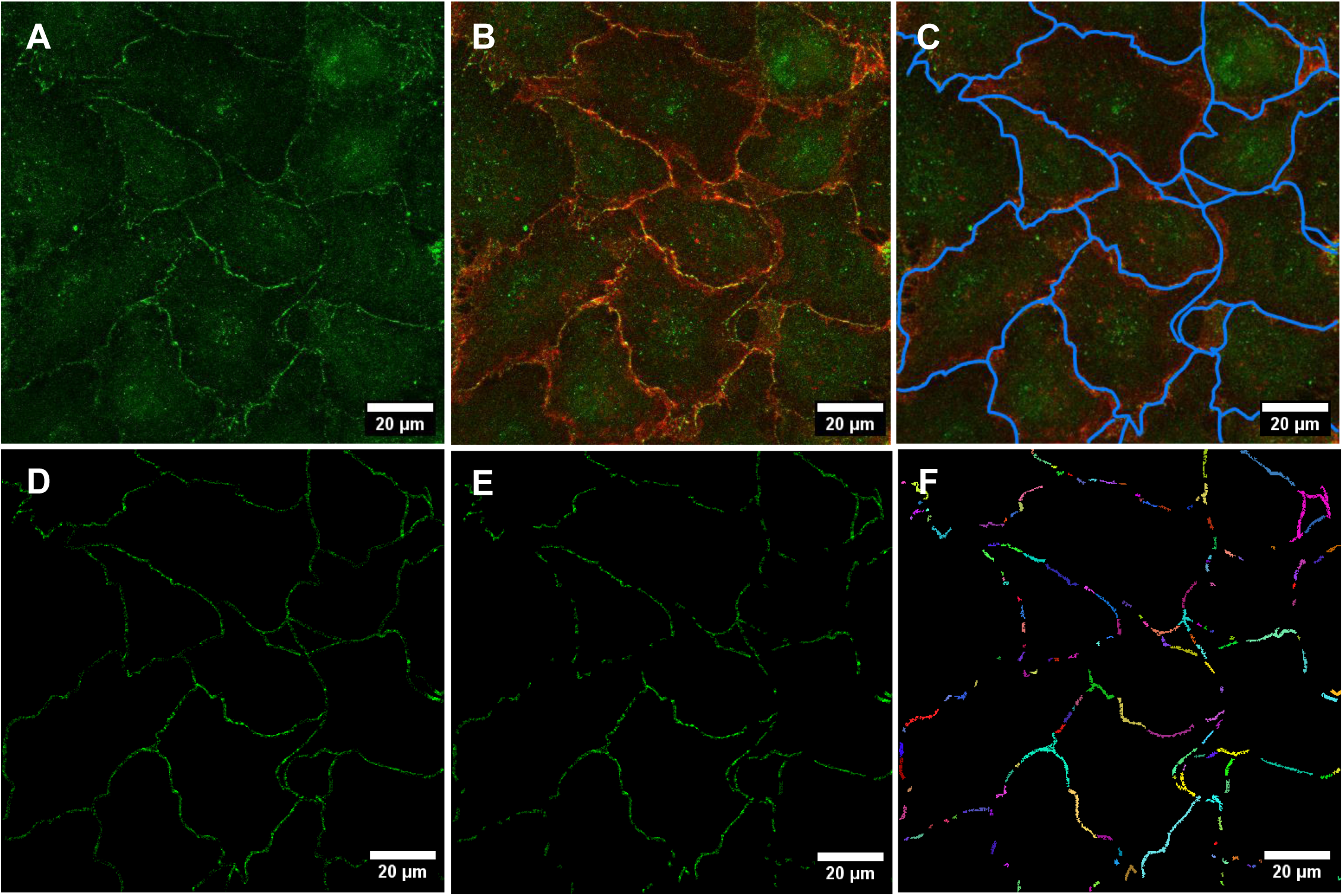
Step-by-step processing of a representative GFP control image through the ZO-1 fragmentation analysis pipeline. **(A)** ZO-1 (green) channel maximum intensity projection showing both junctional signals at cell-cell borders and diffuse cytoplasmic signals throughout the field of view. **(B)** Composite image with VE-cadherin (red) overlaid, identifying cell-cell junctions independently of ZO-1 morphology. **(C)** Manual junction tracings (blue) overlaid on the composite image, guided by VE-cadherin localization. **(D)** ZO-1 signal within the traced junction region after adaptive background correction; signal outside the region and pixels below the image-specific threshold have been removed. **(E)** Junction region after fragment size filtering, in which fragments smaller than 10 pixels have been excluded. **(F)** Individual ZO-1 fragments identified by connected component analysis, each displayed in a unique color. Each fragment is measured for area, perimeter, and compactness. Image acquired on a Zeiss LSM 880 at 63x magnification.

#### 2.2.2 Region of Interest (ROI) Mask Extraction

ZO-1 is present both at cell-cell junctions, where it reflects tight junction integrity, and diffusely in the cytoplasm, where it is not associated with barrier function. To restrict analysis to junctional ZO-1, the junction regions were manually outlined on the exported composite images using blue-colored tracings (**Figure 1C**). Because VE-cadherin localizes exclusively to cell-cell contacts, the red channel served as an independent visual reference for identifying junction location independent of ZO-1 morphology. This approach was analogous to the waypointing step used by Gray et al. in the JAnaP [17], though here the tracing defined a two-dimensional ROI for fragment detection rather than a one-dimensional path for segment classification.

The traced images were imported back into the pipeline using the second notebook, which matched each tracing to its corresponding CZI file by filename. The pipeline automatically detected the blue tracings by isolating pixels in the blue color range. Blue was chosen because it does not overlap with the green (ZO-1) or red (VE-cadherin) channels, allowing the mask to be extracted without interference from the biological signal. Small gaps in the tracings were filled, and isolated noise pixels were removed through standard image cleanup operations. If the traced image differed in pixel dimensions from the original CZI file, the mask was resized to match using nearest-neighbor interpolation. The result was a binary mask defining the junction region.

#### 2.2.3 Automated Fragment Analysis

For each matched CZI-tracing pair, the third notebook loaded the original CZI file, computed the maximum intensity projection, extracted the ZO-1 channel, and applied the corresponding binary mask. All subsequent steps were executed automatically for every image pair in sequence. The complete processing sequence is illustrated for a representative control image in **Figure 1**.

##### 2.2.3.1 Adaptive Background Correction

Applying a single, fixed brightness threshold across all images proved problematic, as background fluorescence varies between imaging sessions and staining batches. A threshold that correctly separated the desired signal from background in one image would over-segment images with brighter backgrounds and under-detect the signal in dimmer images. To address this, the pipeline calculated an image-specific threshold for each input.

For each image, background fluorescence was estimated by calculating the mean fluorescence intensity (MFI) of all pixels outside the manually traced junction ROI. These extra-junctional pixels represented regions where ZO-1 signal was not expected and thus provided a sample of the local background level for that specific image. The background MFI (Ī*_bg_*) was formally defined in **Eq. 1** as:

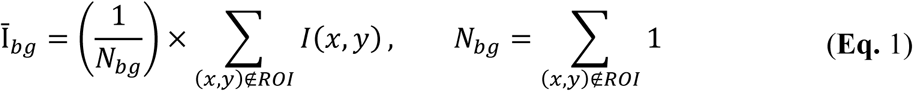

where *I*(*x*, *y*) was the fluorescence intensity of pixel (*x*, *y*), and *N_bg_* was the total number of pixels outside the junction ROI. The adaptive threshold (*T_A_*) was then set as the background MFI (Ī*_bg_*) plus an intensity offset (Δ), as seen in **Eq. 2**:

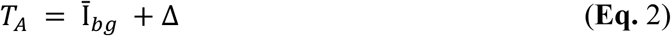

For the *in vitro* analysis, a fixed Δ value of 300 intensity units was empirically determined to effectively separate true ZO-1 junction signal from background fluorescence across all imaging conditions tested in this study. This formulation ensured that the absolute threshold varied with image brightness while maintaining a consistent signal-to-background separation across all images. All pixels within the junction region with intensity below this threshold were excluded from subsequent analysis (**Figure 1D**).

##### 2.2.3.2 Fragment Identification and Morphometric Analysis

After background correction, the remaining bright pixels within the junction region represented ZO-1 signal. Fragments smaller than 10 pixels were excluded to eliminate noise (**Figure 1E**). The pipeline then grouped contiguous bright pixels into discrete fragments: any cluster of touching pixels was identified as a single fragment and assigned a unique label (**Figure 1F**). The pipeline converted pixel areas to physical units automatically using the pixel calibration extracted from the CZI file metadata. At the pixel size used in this study (0.264 μm), the 10-pixel threshold corresponded to 0.71 μm². For each fragment, the following parameters were calculated: area (in μm²), perimeter (the length of the fragment boundary), and compactness (the ratio of fragment area to perimeter, where higher values indicate rounder shapes and lower values indicate elongated or irregular morphologies).

##### 2.2.3.3 Output Metrics and Data Export

Three metrics summarized overall junction integrity per image: (A) average fragment area, where smaller values indicated more severe fragmentation, (B) junctional fragmentation ratio, defined as the fraction of the traced junction region occupied by ZO-1 signal above background, where lower ratios indicated more fragmented junctions, and (C) total junctional area, the summed area of all retained fragments per image.

Per-fragment and per-image metrics were exported to CSV files and organized by experimental condition for downstream statistical analysis. The pipeline generated diagnostic visualizations at each processing stage: the original green channel with the junction region overlay, the brightness-filtered result, and a composite overlay showing retained fragments (green) over the original traced region (red) with the fragmentation ratio annotated.

### 2.3 *IN VIVO* TISSUE PREPARATION AND IMAGING

All animal procedures were conducted under protocols reviewed and approved by the Northeastern University Institutional Animal Care and Use Committee (NU-IACUC), in accordance with the NIH Guide for the Care and Use of Laboratory Animals. Four C57BL/6 mouse cohorts were used: young healthy controls (3 months, n = 4), TBI (3 months, n = 2), aged (18 months, n = 4), and APP/PSEN1dE9 AD model (18 months, n = 4). All mice were female. Female mice in the APPswe/PSEN1dE9 model exhibit more severe amyloid pathology and greater cognitive deficits than age-matched males [47], making them a more stringent test case for detecting AD-associated vascular changes. For the TBI cohort, mild TBI was induced in awake mice using a closed-head momentum exchange impact model [48]. Each TBI mouse received two impacts on consecutive days via a pneumatic impactor fitted with a protective helmet assembly, with impacts centered at -1 mm bregma on the dorsal skull. TBI mice were sacrificed 10 days following the first impact. Together, these cohorts enabled comparative analysis across aging, injury, and amyloid pathology.

#### 2.3.1 Tissue Collection and Processing

Mice were sacrificed by exsanguination, involving transcardial perfusion with phosphate-buffered saline (PBS) containing 0.5% bovine serum albumin (BSA) followed by 4% paraformaldehyde. Brains were subsequently removed, post-fixed overnight in 4% paraformaldehyde at 4°C, cryoprotected in 30% sucrose at 4°C until saturated, and embedded in optimal cutting temperature (OCT) compound at -20°C overnight. All brains were next cryosectioned at 40 μm and collected as free-floating sections for immunohistochemical analysis.

#### 2.3.2 Immunohistochemistry Protocol

Immunohistochemistry was used to visualize specific biomarkers in tissue sections using fluorescently labeled probes. Free-floating sections were first subjected to antigen retrieval by incubation in citrate buffer at 95°C for 20 minutes, followed by three PBS washes. Sections were then blocked and permeabilized in 5% normal goat serum with 0.3% Triton X-100 for one hour. Primary antibodies against the proteins of interest, ZO–1 (rabbit, Invitrogen, catalog 61–7300, 1:100), ICAM–1 (rabbit, Novus Biologicals, catalog NBP2–67518, 1:500), and eNOS (rabbit, Cell Signaling, catalog 32027, 1:100), were applied overnight at 4 °C in a solution of 5% normal goat serum with 0.1% Tween–20 in PBS. Following three PBS washes with 0.1% Tween-20, for secondary detection of these rabbit primary antibodies, Alexa Fluor 555 goat anti–rabbit (Invitrogen, catalog A–21428, 1:500) was applied for two hours at room temperature. DyLight 488-conjugated tomato lectin (Invitrogen, catalog L32470, 1:500) was included for vessel identification. After staining, tissue samples were washed three times before mounting with Vectashield Antifade Mounting Medium with 4′,6–diamidino–2–phenylindole (DAPI; Vector Laboratories), to visualize cell nuclei.

#### 2.3.3 Confocal Imaging of Stained Tissue Sections

Confocal imaging was performed on tissue sections from the PFC, hippocampus, and midbrain areas of the brain. ZO-1 images were acquired at 40x magnification with a 1.4 numerical aperture oil-immersion objective. ICAM-1 images were acquired at 20x magnification with a 0.8 numerical aperture air objective. eNOS images were acquired at either 20x or 40x, depending on the cohort and animal. The image analysis pipeline read magnification from CZI file metadata and adjusted scale calibration accordingly. One representative image per brain region per mouse was acquired for each marker, a sampling strategy chosen to provide a standardized and controlled input set for evaluating the performance of the image–analysis framework rather than to quantify biological variability. These images served as the input dataset for the vascular colocalization pipeline described in Section 2.4.

### 2.4 DEVELOPMENT OF VASCULAR MARKER COLOCALIZATION ANALYSIS PIPELINE

The quantification pipeline described in this section was designed to measure any fluorescent marker within boundaries defined by a second channel. In this study, the reference channel was tomato lectin (defining vessel boundaries) and the markers of interest were ZO-1, ICAM-1, and eNOS, but the same framework applies to any pair of channels where one defines a region and the other is measured within it.

The pipeline operated in three stages. First, a machine learning pixel classifier was trained in ilastik (described in Section 2.4.1) to distinguish blood vessels from microglia in the tomato lectin channel. Second, a Python notebook extracted individual channels from the raw confocal files, applied the trained classifier to generate cleaned vessel images, and prepared the data for analysis. Third, a FIJI macro performed background correction, created a vessel mask, measured protein signal within vessel boundaries, and exported the results. Image files followed a standardized naming convention encoding the protein of interest, mouse cohort, brain region, and sample number. The complete pipeline has been made available as a Jupyter notebook on GitHub (https://github.com/peckbe/ilastik-fiji-colocalization) under the MIT license.

#### 2.4.1 ilastik Pixel Classifier Training

Because the lectin channel contained both vascular and microglial signal in neuroinflammatory cohorts (as discussed in Sections 1.3 and 1.5), and no channel was available for a dedicated microglial counterstain, the pipeline incorporated ilastik, a machine learning tool that uses user-provided examples to classify each pixel in an image [49].

Training proceeded as follows. The tomato lectin channel was extracted from each confocal file as a Z-stack. ilastik requires all input images to have the same number of focal planes, so all Z-stacks were padded with empty slices to match the depth of the deepest stack in the dataset, then exported in HDF5 format for ilastik compatibility. Because the pipeline collapses Z-stacks into maximum intensity projections before analysis, the added empty slices (intensity value zero) cannot override any pixel containing real signal and cannot affect the projected image. Five representative images with high and low microglial content were selected from the young, AD, and TBI datasets. In the ilastik interface, we painted labels on these images to indicate which structures were vessels, which were microglia, and which were background. The corresponding ZO-1 or eNOS channels were displayed alongside the labels as a visual reference to help distinguish the two cell types but were not included as inputs to the classifier. Classification was performed in three dimensions across multiple focal planes rather than on two-dimensional projections, because examining intensity and texture patterns across adjacent planes improved the classifier’s ability to distinguish vessels from microglia, which differ in their Z-axis morphology. Once trained, this classifier was reused for all subsequent images without retraining.

#### 2.4.2 ilastik Prediction and Tomato Lectin Preprocessing

The trained classifier was then applied automatically to all tomato lectin Z-stacks, producing a probability map for each image: at every pixel, the classifier estimated the likelihood that the pixel belonged to a vessel, to a microglial cell, or to background.

These probability maps were used to generate three processed versions of the original tomato lectin image (**Figure 2A**), each representing a different strategy for suppressing microglial signal. The first version (binary mask) assigned each pixel to the class with the highest probability and kept only the vessel-classified pixels (**Figure 2B**). The second version (probability-weighted) scaled the original lectin signal by the vessel probability at each pixel, attenuating signal in regions of low vessel confidence (**Figure 2C**). The third version (dominant-class) provided the most stringent microglial suppression: it examined each focal plane individually and removed slices where microglial probability exceeded vessel probability before collapsing the stack, then further weighted the result by vessel probability (**Figure 2D**).

**Figure 2:**
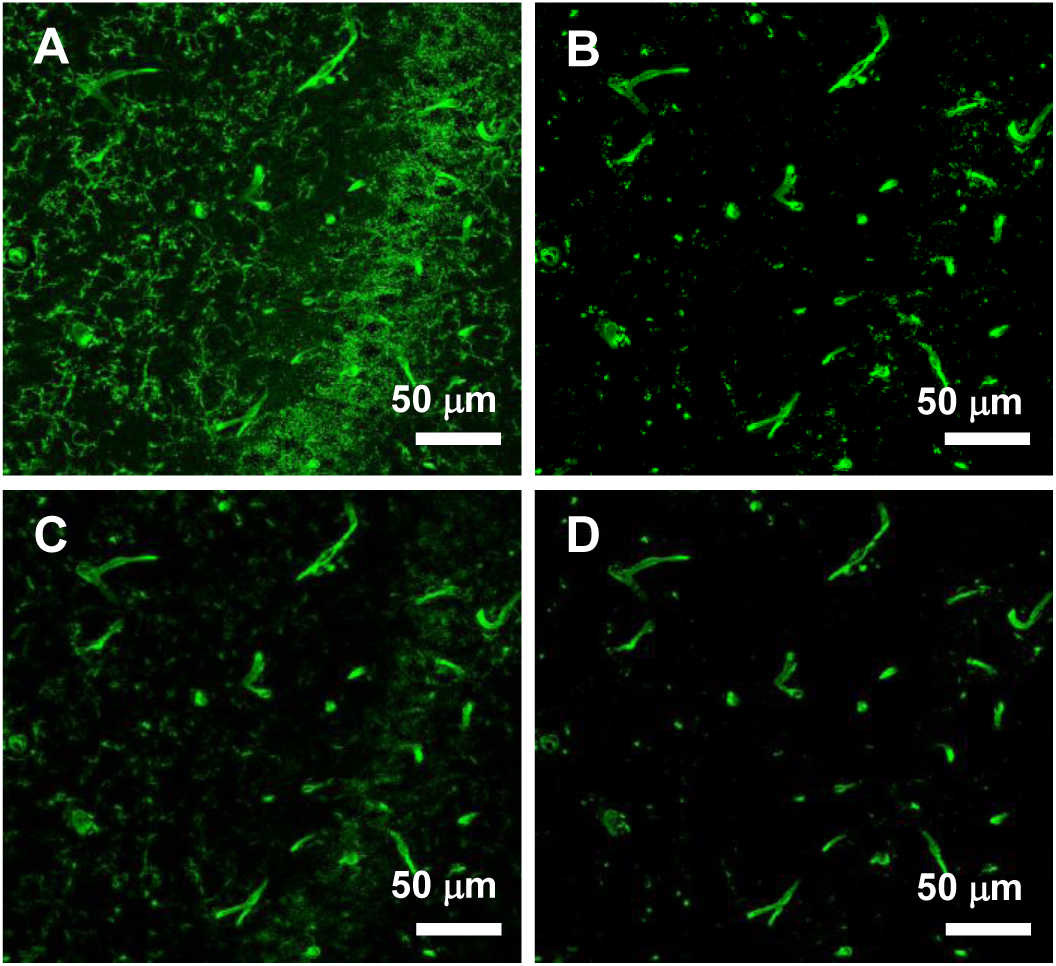
ilastik-based vessel mask variants for a TBI hippocampus image with high microglial content. **(A)** Original tomato lectin maximum intensity projection showing extensive microglial signal throughout the field of view. **(B)** Binary mask, in which each pixel is assigned to the class with the highest probability and only vessel-classified pixels are retained. **(C)** Probability-weighted mask, in which the original lectin signal is scaled by vessel probability at each pixel, attenuating regions of low vessel confidence. **(D)** Dominant-class mask, the most stringent variant, in which focal planes where microglial probability exceeded vessel probability were removed before projection and the result was further weighted by vessel probability. The user selects the variant that most accurately captured vessel boundaries for each image before proceeding with colocalization analysis.

A quality control step displayed all three cleaned versions alongside the original lectin image in a four-panel comparison, allowing the user to visually assess which version best captured the true vessel boundaries before proceeding.

#### 2.4.3 Image Loading and Channel Assignment

For each image, the FIJI macro opened the CZI file via the Bio-Formats plugin (an open-source library for reading proprietary microscopy file formats), which preserved the original metadata and channel structure. Background subtraction was applied first, followed by a maximum intensity projection to collapse the Z-stack into a single two-dimensional image, as in the *in vitro* pipeline. The multi-channel projected image was then split into separate windows for the protein of interest, tomato lectin, and DAPI, allowing each channel to be processed independently.

Channel assignments were determined automatically by the Python notebook, which read fluorophore information stored in the CZI file metadata and mapped each detected fluorophore to its corresponding channel role. When this information was unavailable or ambiguous, the pipeline used pre-configured assignments for each cohort, which could be verified and overridden at runtime.

#### 2.4.4 Background Subtraction

Uneven illumination and tissue autofluorescence can introduce artificial intensity gradients across an image. To correct this, a rolling ball background subtraction [50] with a radius of 50 pixels was applied to all channels across all focal planes before projection.

#### 2.4.5 Reference Channel Mask Generation

The goal of this step was to produce a binary mask from the reference channel: a map of the image where each pixel was classified as either inside or outside the region of interest. For the vascular analysis in this study, the reference channel was tomato lectin, and the mask defined vessel boundaries. All subsequent measurements were restricted to mask-positive pixels only. The approach to generating this mask varied by image, depending on the degree of microglial contamination in the tomato lectin channel.

For images with minimal microglial signal, the tomato lectin channel was thresholded using the Otsu method [51], an algorithm that automatically selects a brightness cutoff to separate foreground from background. The computed threshold was user-verified before it was applied and could be adjusted if the default does not accurately separate the desired signal from the background.

For images with microglial contamination, the pipeline displayed all four tomato lectin options (original plus the three ilastik-processed versions) in a tiled view, and the user selected the version that most accurately captured vessel boundaries. In practice, standard Otsu thresholding was sufficient for images with minimal microglial contamination, while the ilastik-processed versions were selected for images with substantial microglial signal.

Once generated, the resulting binary mask was refined by connecting nearby vessel segments separated by small gaps, filling small internal holes within vessel cross-sections, and removing objects smaller than 15 pixels to exclude noise. The vessel area fraction (the proportion of the image occupied by vessels) was recorded, and the finalized vessel mask was saved so that identical vessel boundaries could be applied if the same image were re-analyzed, ensuring reproducibility across independent analysis runs.

For images where the classifier did not fully resolve residual artifacts, the pipeline supported an optional ROI mode. A freehand ROI was drawn in FIJI to define the analysis region, excluding any artifacts from all downstream measurements. The pipeline reprocessed the image using the same vessel mask and threshold settings but restricted all area calculations to within the ROI boundaries. This was used for a small number of images where retraining the classifier was not justified.

#### 2.4.6 Protein of Interest Colocalization

With the reference channel mask established, the pipeline measured protein signal exclusively within mask boundaries. The protein channel was multiplied by the normalized binary mask, scaled from 0/255 to 0/1. Where the mask was positive, the original protein intensity was preserved. Where the mask indicated negative, the protein signal was set to zero. The MFI of this masked image was recorded as a measure of protein expression intensity within the reference region.

To determine what fraction of the vessel area was positive for the protein, the pipeline used an adaptive threshold related to but distinct from the *in vitro* approach. Background intensity was measured as the median protein signal in non-vessel regions of the image (Ĩ*_bg_*):

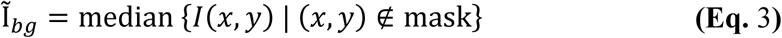

The median was used instead of the mean because non-vessel regions of tissue sections contained intensity outliers in both directions: scattered bright artifacts pulled the mean upward, while dark regions underlying cell nuclei, where membrane-associated proteins were absent, pulled it downward. The median was robust to both sources of skew, providing a more stable estimate of true background fluorescence. The threshold (*T_A_*) was set as:

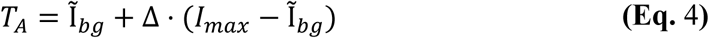

where *I_max_* is the maximum protein intensity within the image and Δ = 0.15 for all markers.

The adaptive threshold was then applied to the multiplied image, converting it to a binary mask of protein-positive pixels within the reference region. Protein coverage was calculated as the ratio of the protein-positive area (measured as percent area of the full image) to the reference channel area (also measured as percent area of the full image), expressed as a percentage. Because the coverage metric normalizes protein-positive area to vessel area, it is independent of differences in vessel density between images. The vessel area fraction was recorded for each image but was not used as an outcome variable.

#### 2.4.7 Output and Organization

Results were exported to a CSV file containing reference channel area fraction, protein MFI, protein coverage, and percent protein coverage within the reference region for each image. The pipeline automatically organized output directories by mouse cohort, brain region, and protein. For each image, the pipeline saved four diagnostic images automatically: the reference channel mask, the protein channel, a composite overlay showing the reference channel in grayscale with thresholded protein signal in red, and the colocalization binary mask. All saved images included scale bars sized to the objective magnification. Reference channel masks were saved to a shared directory so that identical boundaries could be reloaded if the same image was re-analyzed.

#### 2.4.8 Blinding

To minimize potential bias during user-dependent analysis steps, the pipeline incorporated a blinding protocol. Before processing, images were randomized and each was assigned a generic identifier (e.g., Image_001). All FIJI analysis windows displayed generic channel labels (TL, Protein, Overlay) rather than filenames, and the FIJI log was cleared at the start of each image to prevent accidental exposure of file metadata. Dialog prompts for ROI selection, vessel mask thresholding, and ilastik method selection displayed only the generic identifier without cohort, brain region, or protein information. An unblinding key that mapped identifiers back to the original filenames was generated after all images in the batch were processed.

### 2.5 STATISTICAL ANALYSIS

Statistical analyses were performed using GraphPad Prism. For the *in vitro* ZO-1 fragmentation data, per-image metrics were averaged within each independent flow experiment to yield one value per experiment per condition. Comparisons across the three experimental conditions (GFP control, n = 5; CD44 KD, n = 4; SDC1 KD, n = 5) were conducted using ordinary one-way ANOVA with Tukey’s post-hoc multiple comparisons test. Homogeneity of variance was confirmed by Brown-Forsythe and Bartlett’s tests, which verify that variance is comparable across groups, a requirement for valid ANOVA results.

For the *in vivo* vascular marker data, a two-way ANOVA (cohort x brain region) with Tukey’s post-hoc multiple comparisons was performed separately for each marker (ZO-1, ICAM-1, eNOS). TBI mice were included as a positive control for BBB disruption and are presented descriptively in all figures but were excluded from the statistical model due to reduced sample size (n = 2 vs. n = 4 for the experimental groups). The formal statistical comparisons were therefore restricted to three groups: young (n = 4), aged (n = 4), and AD (n = 4). Two-way ANOVAs were performed on the three regional measurements (PFC, hippocampus, midbrain). Global averages, computed by averaging the three regional values for each mouse, were analyzed separately using one-way ANOVA with Tukey’s post-hoc correction. Data are presented as mean ± SEM. Statistical significance was set at p < 0.05.

## 3. RESULTS AND DISCUSSION

### 3.1 *IN VITRO* ZO-1 FRAGMENTATION ANALYSIS

#### 3.1.1 Results

The ZO-1 fragmentation pipeline was applied to confocal images of HBMECs under three experimental conditions: GFP control, CD44 KD, and SDC1 KD. Representative images illustrate the transition from near-continuous, well-defined ZO-1 staining at cell-cell junctions in GFP control cells (**Figure 3A**) to the dim, discontinuous junctions in CD44 KD (**Figure 3B**) and SDC1 KD cells (**Figure 3C**). Pipeline overlays showing detected fragments are presented for each image (**Figure 3D-F**).

**Figure 3:**
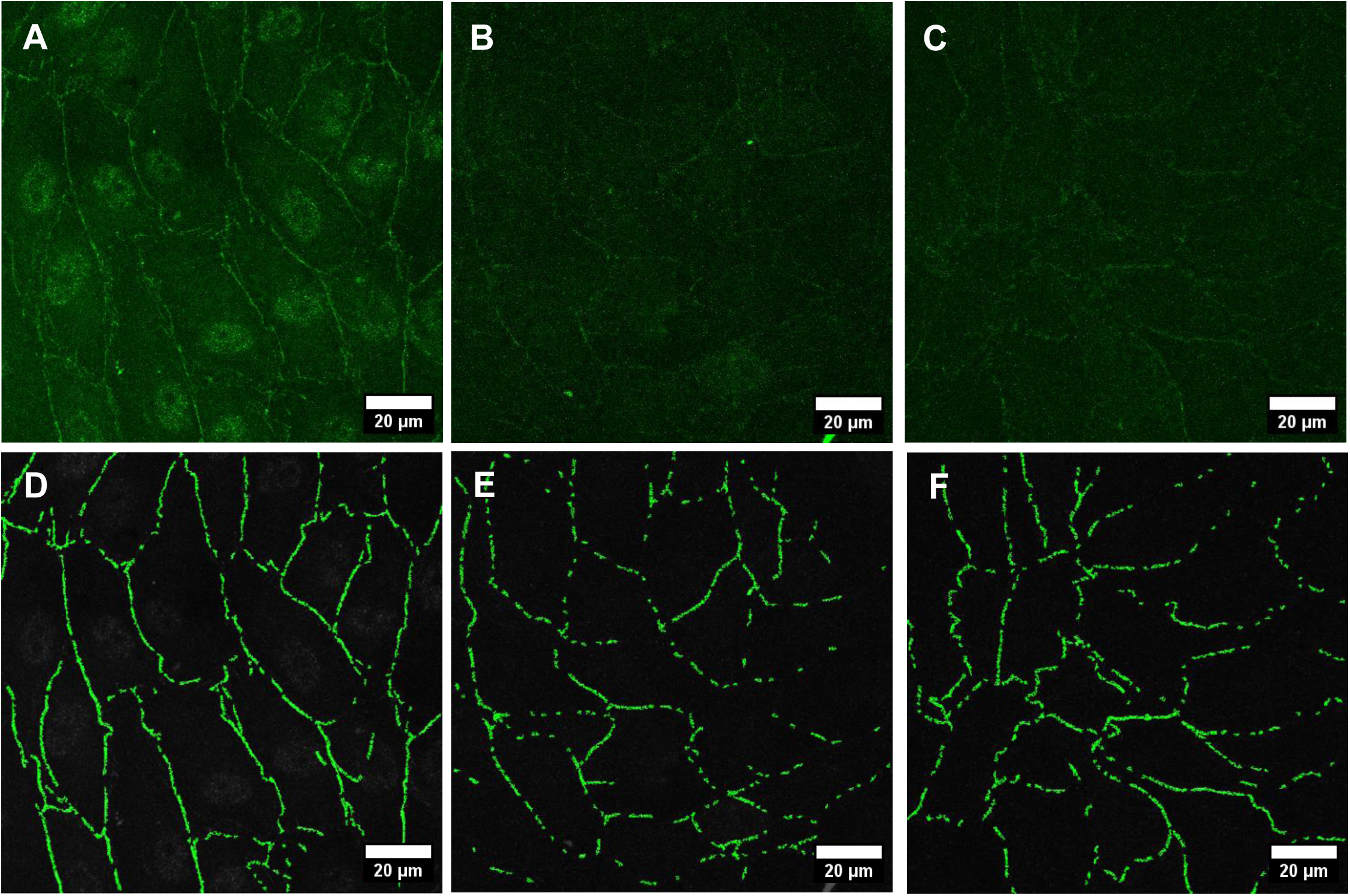
ZO-1 junction morphology is disrupted in GCX KD conditions, as detected by the fragmentation analysis pipeline. **(A-C)** Representative confocal images of ZO-1 (green) in GFP control **(A)**, CD44 KD **(B)**, and SDC1 KD **(C)** HBMECs. Control cells exhibit continuous ZO-1 bands at cell-cell junctions, while both CD44 and SDC1 KD conditions show fragmented, discontinuous staining. **(D-F)** Corresponding pipeline analysis showing retained ZO-1 fragments (green) within the traced junction region for GFP control **(D)**, CD44 KD **(E)**, and SDC1 KD **(F)**. Non-junctional signal has been removed, isolating only fragments within the traced region. Images were acquired on a Zeiss LSM 880 at 63x magnification.

Three metrics captured the fragmentation phenotype, each displayed in **Figure 4** normalized to the GFP control and reported in the text as absolute values. Average fragment area (**Figure 4A**), decreased from 2.84 μm² in GFP controls to 1.80 μm² in CD44 KD (p = 0.0148, **Figure 4D**) and 1.76 μm² in SDC1 KD (p = 0.0084, **Figure 4D**), representing an approximately 37% reduction for both KD conditions. CD44 KD and SDC1 KD did not differ from each other (p = 0.9925, **Figure 4D**). The junctional fragmentation ratio (**Figure 4B**), defined as the proportion of the traced junction region occupied by ZO-1 signal above background, decreased from 43.44% in controls to 26.85% in CD44 KD (p = 0.0061, **Figure 4D**) and 29.68% in SDC1 KD (p = 0.0137, **Figure 4D**), with no difference between KD conditions (p = 0.7852, **Figure 4D**).

**Figure 4:**
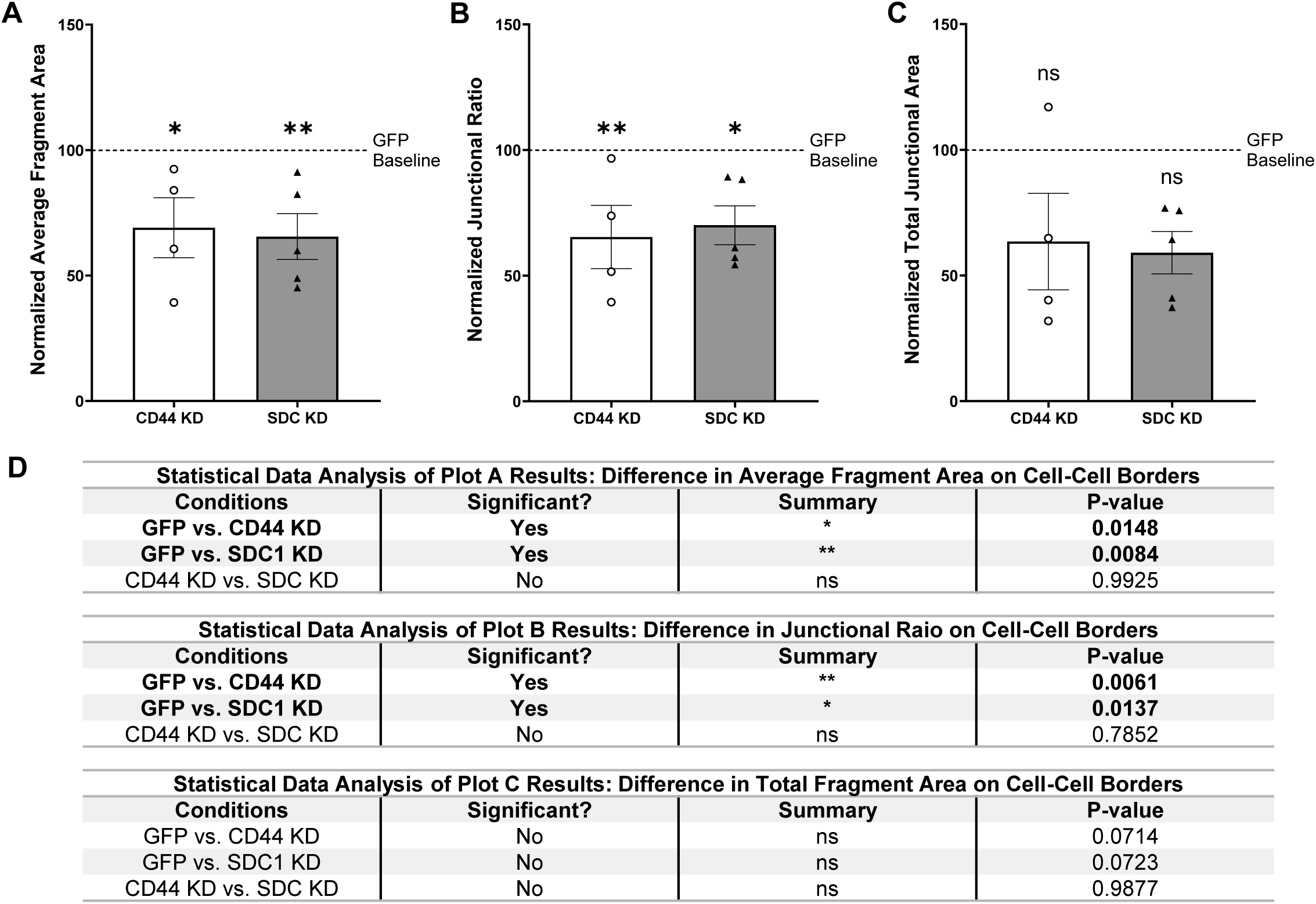
ZO-1 fragmentation metrics are significantly reduced in both GCX KD conditions relative to control. **(A)** Average fragment area, reflecting the mean size of individual ZO-1 fragments per image, normalized to the GFP control group (GFP = 100%, dashed line). **(B)** Junctional fragmentation ratio, the proportion of the traced junction region occupied by ZO-1 signal above background, normalized to the GFP control group (GFP = 100%, dashed line). **(C)** Total junctional area, the summed area of all retained fragments per image, normalized to the GFP control group (GFP = 100%, dashed line). **(D)** Statistical analysis results for the datasets presented in A–C, including group comparisons and corresponding p–values. **(A-D)** Data are presented as mean ± SEM; n = 5 independent flow experiments per condition, except CD44 KD (n = 4). The overall ANOVA detected a significant effect, but no individual pairwise comparison was significant after Tukey’s correction. CD44 KD and SDC1 KD were statistically indistinguishable across all three metrics. *p < 0.05, **p < 0.01, ns = not significant; one-way ANOVA with Tukey’s post-hoc test.

Total junctional area (**Figure 4C**), defined as the summed area of all retained fragments per image, trended toward reduction: 598.44 μm² in controls versus 321.01 μm² in CD44 KD and 337.69 μm² in SDC1 KD. The overall ANOVA detected a significant effect across groups (F_2,11_ = 4.200, p = 0.0441), but no individual pairwise comparison reached significance after Tukey’s correction (Control vs CD44 KD: p = 0.0714, Control vs SDC1 KD: p = 0.0723, CD44 KD vs SDC1 KD: p = 0.9877; **Figure 4D**). The approximately 45% reduction did not reach pairwise significance but follows the same pattern as the fragment area and junctional fragmentation ratio results, likely reflecting smaller fragments occupying less total area within the junction region.

#### 3.1.2 Discussion

The fragmentation results are as expected given our previous observation that adverse conditions disrupt tight junction organization and concurrently erode glycocalyx structure [21]. The significantly reduced fragment area and junctional fragmentation ratio provide reproducible, numerical measures of that disruption. The trend toward reduced total junctional area follows the same pattern and suggests overall ZO-1 loss, though larger sample sizes would be needed to confirm this at the pairwise level. CD44 and SDC1 KD produced nearly identical fragmentation profiles across all three metrics, demonstrating that the pipeline resolves a consistent phenotype regardless of which upstream perturbation caused the disruption. The biological interpretation of these GCX-dependent effects is the focus of ongoing work investigating how specific GCX components regulate tight–junction organization in relation to endothelial barrier integrity. .

The fragment-level metrics go beyond what categorical approaches can capture. The JAnaP, for instance, classifies junction signal into bins along a one-dimensional path (Section 1.5; [17]). The fragmentation pipeline instead measures discrete two-dimensional fragment objects, resolving both individual fragment size and the overall proportion of the junction region occupied by ZO-1. The pipeline also uses an adaptive threshold (**Eq. 2**) that accounts for image-to-image variation in background intensity, avoiding the systematic bias that fixed-threshold approaches introduce across conditions with different staining characteristics.

### 3.2 *IN VIVO* VASCULAR MARKER ANALYSIS

#### 3.2.1 Results

The *in vivo* analysis shifted our focus beyond controlled cell culture to tissue from whole-animal disease models, where age, genetic background, and injury history vary simultaneously. Each marker was quantified independently within vessel boundaries defined by the tomato lectin channel.

##### 3.2.1.1 ZO-1: Tight Junction Integrity

ZO-1 percent area coverage on vessels was quantified across three brain regions in young, aged, and AD cohorts using the colocalization pipeline. Representative confocal images from the PFC illustrate ZO-1 signal on lectin-labeled vessels in young controls (**Figure 5A**) and show the reduction in ZO-1 signal in aged (**Figure 5B**) and AD mice (**Figure 5C**), with TBI mice showing the most severe reduction (**Figure 5D**).

**Figure 5:**
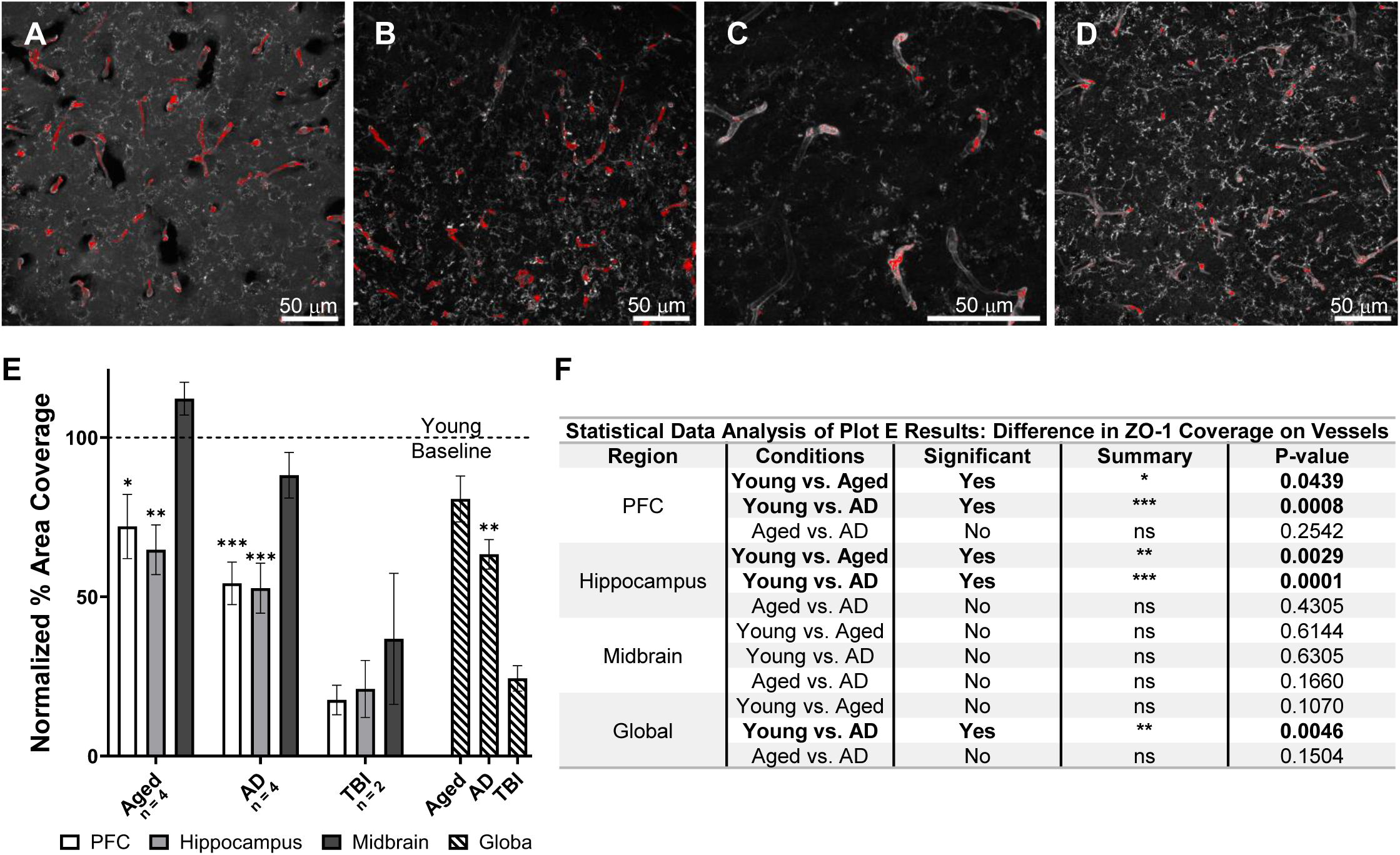
ZO-1 coverage on vessels decreases in aged and AD mice, with the hippocampus and PFC most affected. **(A-D)** Representative confocal images from the PFC showing tomato lectin (gray) and ZO-1 (red) in young **(A)**, aged **(B)**, AD **(C)**, and TBI **(D)** mice. All images were acquired at 40x magnification, and the young, aged and TBI images are tile scans encompassing a larger field of view. **(E)** Quantification of ZO-1 percent area coverage on vessels in the PFC, hippocampus, and midbrain, and as a global average across regions, normalized to the young group within each region (young = 100%, dashed line). TBI data are shown descriptively and were excluded from the statistical model. **(F)** Statistical analysis results for the datasets presented in **E**, including group comparisons and corresponding p–values. **(E, F)** Data are presented as mean ± SEM; n = 4 per group except TBI (n = 2). Regional comparisons: two-way ANOVA with Tukey’s post-hoc test. Global comparisons: one-way ANOVA with Tukey’s post-hoc test. Significance is denoted as *p < 0.05, **p < 0.01, ***p < 0.001, ns = not significant.

The image analysis pipeline reveals clear region–specific differences in tight–junction coverage in young mice, with significantly higher ZO-1 coverage in the hippocampus than in the midbrain (67.40% vs 50.28%, p = 0.0343). Although regional variation in BBB properties has been previously recognized at the transcriptomic level [52, 53] and at the functional level, where BBB permeability is higher in the cortex and hippocampus than in the midbrain [54], to our knowledge, this is the first quantification of ZO-1 protein differences in adult C57BL/6 mice.

Extending this analysis to aging and disease, the hippocampus showed the most pronounced reductions in ZO-1 coverage (**Figure 5E**). Compared to young mice, which displayed the highest coverage at 67.40%, ZO-1 decreased to 43.64% in aged mice (p = 0.0029, **Figure 5F**) and 35.52% in AD mice (p = 0.0001, **Figure 5F**). Aged and AD mice were not significantly different from each other despite the numerically lower AD average (p = 0.4305, **Figure 5F**). In the PFC, a similar pattern was observed: ZO-1 coverage decreased from 58.69% in young mice to 42.30% in aged mice (p = 0.0439, **Figure 5F**) and 31.83% in AD mice (p = 0.0008), with no significant difference between aged and AD conditions (p = 0.2542, **Figure 5F**). The midbrain showed a distinct pattern: coverage was statistically similar in young mice (50.28%), aged (56.41%), and AD mice (44.32%), with none of the pairwise comparisons reaching significance (all p > 0.16, **Figure 5F**). TBI mice showed the lowest ZO-1 coverage of any cohort across all three regions, consistent with established acute tight junction loss following injury [31].

Globally, young mice exhibited higher ZO-1 coverage (58.79%) than both aged (47.45%) and AD mice (37.22%). Young and AD mice differed significantly (p = 0.0046, **Figure 5F**), while the young-to-aged (p = 0.1070, **Figure 5F**) and aged-to-AD (p = 0.1504, **Figure 5F**) comparisons did not reach significance.

##### 3.2.1.2 ICAM-1: Endothelial Activation

ICAM-1 percent area coverage on vessels was quantified across the same three brain regions and cohorts. Representative images from the PFC show that, compared to young controls (**Figure 6A**), ICAM-1 signal increased on the vasculature in aged mice (**Figure 6B**) and AD mice (**Figure 6C**). TBI mice (**Figure 6D**) showed coverage that had largely returned to young control levels except in the hippocampus, directly underlying the impact site.

**Figure 6:**
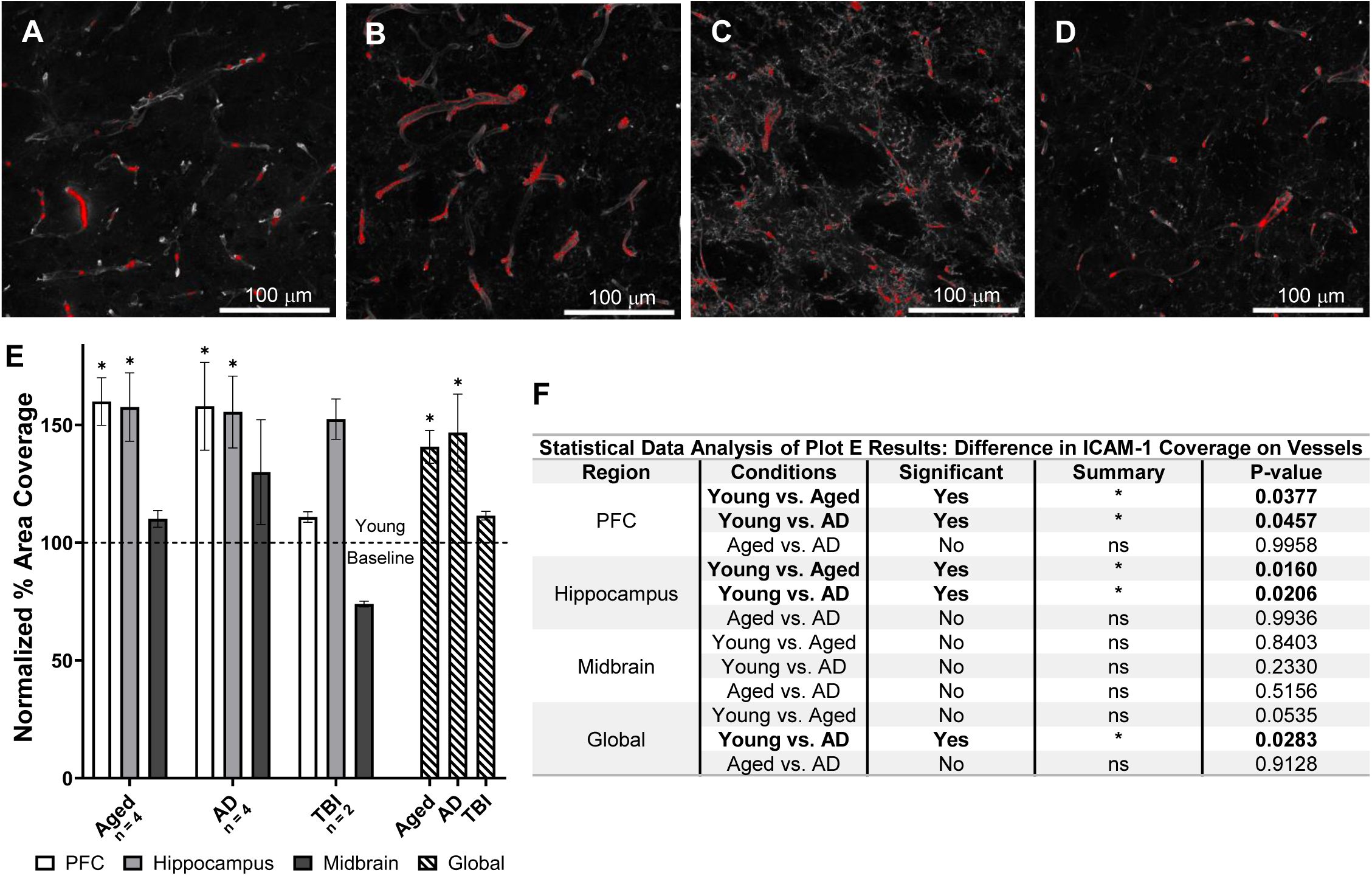
ICAM-1 coverage on vessels increases in aged and AD mice, with the hippocampus and PFC most affected. **(A-D)** Representative confocal images from the PFC showing tomato lectin (gray) and ICAM-1 (red) in young **(A)**, aged **(B)**, AD **(C)**, and TBI **(D)** mice. All images were acquired at 20x magnification. **(E)** Quantification of ICAM-1 percent area coverage on vessels in the PFC, hippocampus, and midbrain, and as a global average across regions, normalized to the young group within each region (young = 100%, dashed line). TBI data are shown descriptively and were excluded from the statistical model. **(F)** Statistical analysis results for the datasets presented in **E**, including group comparisons and corresponding p–values. **(E, F)** Data are presented as mean ± SEM; n = 4 per group except TBI (n = 2). Regional comparisons: two-way ANOVA with Tukey’s post-hoc test. Global comparisons: one-way ANOVA with Tukey’s post-hoc test. Significance is denoted as *p < 0.05, ns = not significant.

In the PFC, ICAM-1 coverage increased from 34.30% in young mice to 54.86% in aged mice (p = 0.0377, **Figure 6F**) and 54.17% in AD mice (p = 0.0457, **Figure 6F**), with aged and AD groups virtually identical (**Figures 6E** and **6F**). The hippocampus showed the most pronounced elevation, with coverage rising from 40.81% in young mice to 64.30% in aged mice (p = 0.0160, **Figure 6F**) and 63.45% in AD mice (p = 0.0206, **Figure 6F**). Aged and AD were virtually indistinguishable (**Figures 6E** and **6F**). In the midbrain, coverage ranged from 44.06% in young mice, to 48.51% in aged mice and 57.26% in AD mice, but none of these differences reached significance (all p > 0.23, **Figures 6E** and **6F**).

TBI mice showed ICAM-1 coverage that had largely returned to young control levels in the PFC and midbrain, consistent with the transient nature of post-traumatic ICAM-1 upregulation at the 10-day sacrifice timepoint [32]. The hippocampus was an exception, where TBI mice showed ICAM-1 coverage comparable to the aged and AD cohorts.

Globally, young mice exhibited lower ICAM-1 coverage (39.73%) than both aged (55.89%) and AD mice (58.29%). Young mice differed significantly from AD mice (p = 0.0283, **Figure 6F**) but not from aged mice (p = 0.0535, **Figure 6F**), while aged and AD mice were not significantly different (**Figures 6E** and **6F**).

##### 3.2.1.3 eNOS: Nitric Oxide Mediated Endothelial Signaling, Barrier Regulation, and Vascular Tone

eNOS percent area coverage on vessels was quantified across the same three brain regions and cohorts. Representative images from the PFC illustrate eNOS signal on vessels in young controls (**Figure 7A**), aged mice (**Figure 7B**), AD mice (**Figure 7C**), and TBI mice (**Figure 7D**). Relative to young mice, AD mice show the greatest increase in eNOS levels, while aged and TBI mice exhibit more moderate increases.

**Figure 7:**
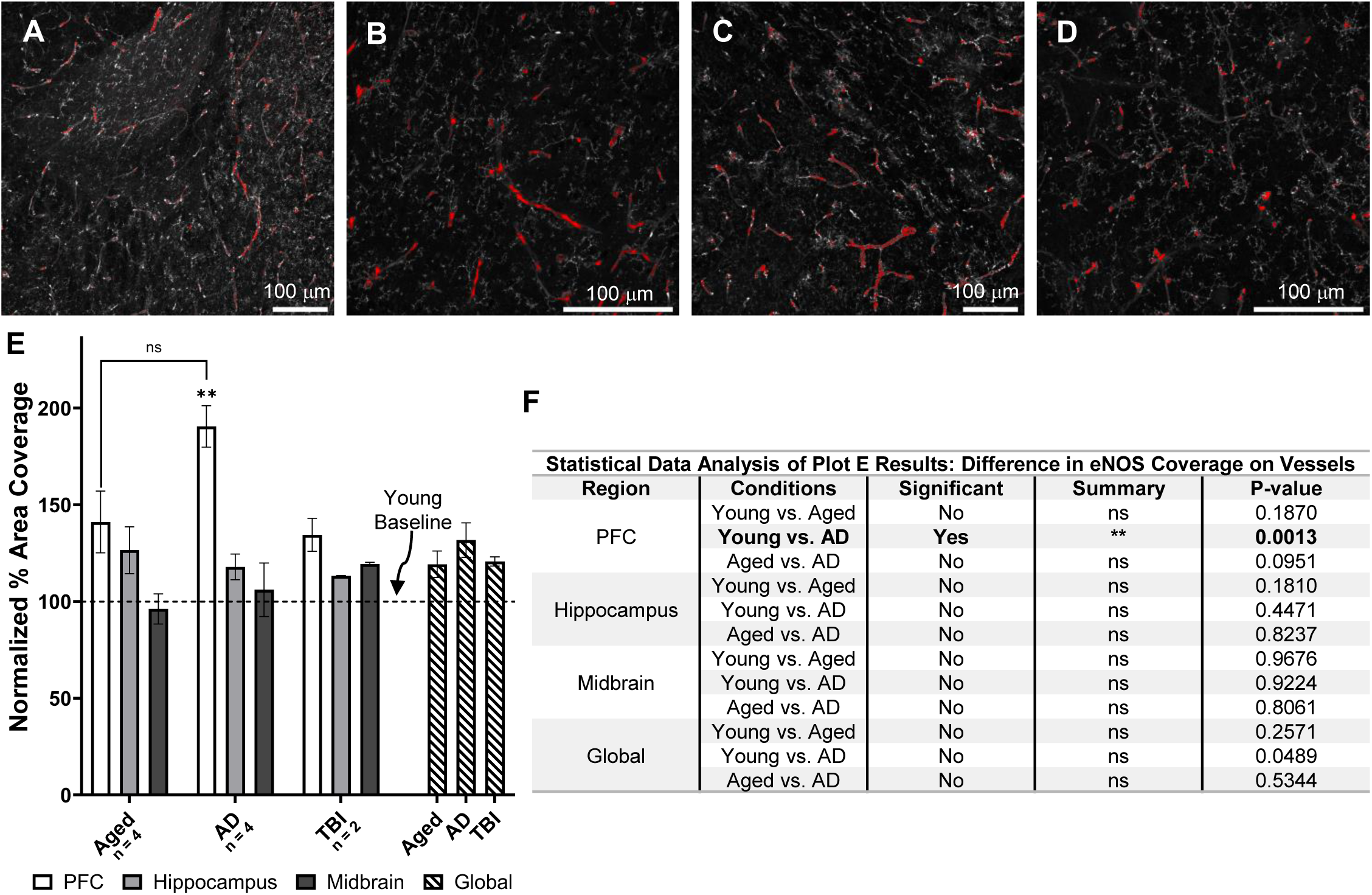
eNOS coverage on vessels increases selectively in the PFC of AD mice. **(A-D)** Representative confocal images from the PFC showing tomato lectin (gray) and eNOS (red) in young **(A)**, aged **(B)**, AD **(C)**, and TBI **(D)** mice. Young and AD images were acquired at 20x, while aged and TBI images were acquired at 40x magnification. Young and AD images are tile scans encompassing a larger field of view. **(E)** Quantification of total eNOS percent area coverage on vessels in the PFC, hippocampus, and midbrain, and as a global average across regions, normalized to the young group within each region (young = 100%, dashed line). TBI data are shown descriptively and were excluded from the statistical model. **(F)** Statistical analysis results for the datasets presented in **E**, including group comparisons and corresponding p–values. **(E, F)** Data are presented as mean ± SEM; n = 4 per group except TBI (n = 2). Regional comparisons: two-way ANOVA with Tukey’s post-hoc test. Global comparisons: one-way ANOVA with Tukey’s post-hoc test, however, since the global one-way ANOVA was nonsignificant (p = 0.057), the post-hoc comparisons were not interpreted. Significance is denoted as **p < 0.01, ns = not significant.

The PFC was the only region to yield a significant result, with **Figures 7E** highlighting that eNOS coverage increased from 33.20% in young mice to 63.25% in AD mice (p = 0.0013, **Figure 7F**). Young and aged mice did not differ significantly (46.85%, p = 0.1870, **Figures 7E** and **7F**), nor did aged and AD mice (p = 0.0951, **Figure 7F**). In the hippocampus, coverage ranged from 52.02% in young to 65.82% in aged mice and 61.31% in AD mice, but none of these differences were significant (all p > 0.18, **Figure 7F**). The midbrain showed no significant differences, with means ranging from 45.97% to 50.72% (**Figure 7E**). TBI mice showed eNOS coverage comparable to the aged cohort across all regions, consistent with the resolution of transient post-traumatic eNOS upregulation by the 10-day sacrifice timepoint [33, 34].

Globally, young mice exhibited lower eNOS coverage (44.35%) than both aged (52.88%) and AD mice (58.43%), as shown in **Figure 7E**. The overall one-way ANOVA was not significant (p = 0.057), so none of the pairwise comparisons are interpreted as significant (young vs AD p = 0.0489, young vs aged p = 0.2571, aged vs AD p = 0.5344); see **Figure 7F**.

##### 3.2.1.4 Integrated Vascular Marker Profile

The three vascular markers reveal a consistent regional pattern of BBB-associated changes across cohorts. ZO-1 coverage decreased, and ICAM-1 coverage increased in the PFC and hippocampus of aged and AD mice relative to young controls, indicating concurrent tight junction loss and endothelial activation in these forebrain regions. eNOS coverage increased significantly in the PFC, where AD mice differed from young controls (p = 0.0013). The aged-to-AD prefrontal comparison did not reach significance (p = 0.0951). The midbrain showed no significant differences among groups for any of the three markers. Across all regions and markers, aged and AD mice were not statistically distinguishable, suggesting that the vascular changes detected by the pipeline in the APPswe/PS1dE9 model at 18 months are largely age-driven.

TBI mice showed the expected pattern of acute BBB disruption across all three markers. ZO-1 coverage was markedly reduced across all regions, ICAM-1 showed coverage that had largely returned to baseline except in the hippocampus, and eNOS showed levels comparable to those of the aged cohort. This pattern matches published recovery timelines and shows that distinct marker profiles are detectable within a single cohort.

#### 3.2.2 Discussion

##### 3.2.2.1 ZO-1 and ICAM-1: Aging as the Primary Driver of Vascular Changes

The most consistent finding across the *in vivo* analysis was that aged and AD mice were statistically indistinguishable for both ZO-1 and ICAM-1 in every brain region examined. Both markers changed significantly relative to young controls in the PFC and hippocampus (ZO-1 coverage decreased while ICAM-1 coverage increased), but these changes tracked with aging rather than AD-specific pathology. This was unexpected given existing evidence from human studies that AD accelerates BBB breakdown beyond normal aging [4, 6, 55], but is consistent with the limited direct evidence in this model. Vellonen et al. reported similar tight junction protein profiles between APdE9 and age-matched wild-type control mice at 12-16 months [56], and Nozohouri et al. found no significant differences in ZO-1, claudin-5, or occludin in the Tg2576 AD mouse model [57]. As described in Section 1.4, the APPswe/PSEN1dE9 model produces predominantly Aβ42 and has limited cerebral amyloid angiopathy, potentially explaining why direct endothelial effects are limited compared to models with greater vascular amyloid burden [8].

These findings are consistent with the age-dependent vascular component of AD emphasized by the vascular hypothesis [6], though they do not distinguish whether the observed changes contribute to disease progression or are a consequence of it. The observation that vascular marker changes in AD mice were indistinguishable from normal aging suggests that BBB dysfunction at this time point reflects the age-dependent vascular component of AD pathology rather than a disease-specific insult, though the study was not powered to detect moderate AD-specific effects (see Section 4.2). The concurrent ICAM-1 elevation is also consistent with age-related GCX degradation, as heparan sulfate loss has been shown to directly upregulate ICAM-1 and other adhesion molecules in endothelial cells independent of flow conditions [58].

The regional distribution of marker changes supports this interpretation. The hippocampus, which is known to be susceptible to age-related vascular dysfunction [35, 36], showed the strongest effects for both markers. The PFC showed a similar but slightly attenuated pattern, in line with recent evidence identifying the PFC as the most vulnerable region to age-related BBB changes in mice [59]. The midbrain was spared, mirroring the gradient of age-related BBB vulnerability rather than the cortical distribution of amyloid pathology in this model [37].

TBI mice provided qualitative validation of expected marker behavior. ZO-1 coverage was markedly reduced across all regions, well below even the AD cohort, confirming that tight junction disruption persists at 10 days post-injury. This is consistent with reports that structural BBB damage following TBI can outlast the acute inflammatory response, with tight junction protein loss including ZO-1 documented days to weeks after injury [29, 31]. ZO-1 loss persisted across all three regions, including the midbrain, which was otherwise unaffected in the aging and AD cohorts, indicating that even mild repetitive head trauma produces substantial barrier damage.

ICAM-1 showed a contrasting temporal pattern. Coverage had largely returned to young control levels in the PFC and midbrain, consistent with the known time course of post-traumatic ICAM-1 expression, which peaks within days of injury and returns to baseline within approximately one week [32, 60], placing the 10-day sacrifice timepoint beyond the expected window of elevated expression in most regions. The hippocampus was an exception, where ICAM-1 remained elevated at levels comparable to aged and AD mice, possibly reflecting persistent inflammation directly underlying the dorsal impact site [48]. The dissociation between ZO-1 (still depressed in all regions) and ICAM-1 (resolved except at the impact site) at day 10 illustrates the distinct recovery timelines of structural versus inflammatory components of post-traumatic BBB disruption [28, 30].

##### 3.2.2.2 eNOS: AD-Specific Upregulation in the PFC

eNOS was the only marker to show an AD-specific effect beyond normal aging. In the PFC, eNOS coverage increased from 33.20% to 63.25% in AD mice relative to young controls (p = 0.0013), while aged mice did not differ significantly from either group. This pattern contrasts with ZO-1 and ICAM-1, where aged and AD mice changed in parallel, and suggests that the PFC vasculature exhibits a response to AD pathology that is distinct from the age-related changes captured by the other two markers. Partial eNOS deficiency has been shown to exacerbate both cognitive decline and amyloid pathology in this model [61], further supporting a functional role for eNOS dysregulation in AD-associated vascular changes.

This finding is consistent with Santhanam et al., who reported increased eNOS protein expression in the cerebral vasculature of Tg2576 AD mice alongside markers of oxidative stress [62]. The interpretation is that chronic oxidative stress in AD triggers compensatory upregulation of eNOS protein, but the additional enzyme is dysfunctional. This phenomenon, known as eNOS uncoupling, causes the enzyme to produce harmful reactive oxygen species instead of NO due to loss of essential cofactors [27]. Because eNOS-derived NO normally helps maintain barrier integrity by stabilizing VE-cadherin at endothelial junctions [24, 26], the loss of functional NO in uncoupled eNOS, combined with increased oxidative stress [27], may shift the endothelium from a barrier-protective to a barrier-damaging state. If this process occurs in the APPswe/PSEN1dE9 PFC, it could contribute to the concurrent ZO-1 loss and ICAM-1 elevation observed in the same region: loss of NO-dependent barrier maintenance and anti-inflammatory signaling could contribute to both tight junction disruption and endothelial activation. The PFC may be uniquely susceptible to this response because it continues to accumulate amyloid plaques beyond 12 months in APPswe/PSEN1dE9 mice, when other cortical regions have largely plateaued [63]. The hippocampus, by contrast, showed no significant eNOS changes across any group. Whether this reflects regional differences in baseline eNOS expression or oxidative stress buffering capacity [36] cannot be determined from the present data.

TBI mice showed eNOS coverage that did not differ significantly from either young or aged mice across all regions, contrasting with the persistent ZO-1 loss and the region-dependent ICAM-1 pattern observed in the same animals. Post-traumatic eNOS upregulation peaks within 24 to 72 hours and resolves by approximately six days [33, 34]. The return to near-baseline levels by the 10-day sacrifice timepoint is consistent with this established timeline and further illustrates the marker-specific recovery dynamics following acute injury: ZO-1 loss persists across all regions, ICAM-1 resolves in all regions except the hippocampus (near the impact site), and eNOS returns to aged-cohort levels.

##### 3.2.2.3 Vessel-Microglia Classification: A Novel Analytical Contribution

No existing tool distinguishes two cell types labeled by the same fluorescent marker within a single channel. The ilastik-based classification used here solves this for vessels and microglia in the lectin channel. Prior machine learning approaches to microglial and vascular segmentation rely on dedicated channels where only one cell type is present (Section 1.5). The approach taken here operates entirely within the lectin channel, using morphological and textural features across focal planes to distinguish vessels from microglia without requiring an additional stain. Without this step, microglial contamination of the vessel mask in the TBI and AD cohorts would have inflated vessel area measurements and diluted the colocalization signal for all three markers. Other pixel classification tools, such as WEKA Trainable Segmentation in FIJI, could perform a similar role, however ilastik was chosen for its native three-dimensional Z-stack classification and its batch prediction workflow, both of which were important for exploiting morphological differences between vessels and microglia across focal planes and integrating the classification step into the Python preprocessing notebook.

## 4. LIMITATIONS AND FUTURE DIRECTIONS

### 4.1 *IN VITRO* PIPELINE LIMITATIONS

The *in vitro* pipeline retains a manual junction tracing step. While this step relies on the user’s ability to distinguish junctional from cytoplasmic ZO-1 using VE-cadherin as a biological reference, it introduces subjectivity at the ROI definition stage and limits throughput when large image sets are processed. The tracing also requires VE-cadherin co-staining, which adds an experimental constraint that may not be feasible in all staining panels.

The pipeline operates on two-dimensional maximum intensity projections, collapsing Z-stack data along the axial dimension before analysis. This may underestimate fragment size or overcount fragments when a single three-dimensional junction structure spans multiple Z-planes and appears as separate objects in the projection. Fragment area and perimeter are therefore two-dimensional approximations of three-dimensional structures.

The adaptive threshold uses a fixed intensity offset added to the per-image background average. While the adaptive framework accounts for image-to-image variation in absolute intensity, the offset itself was empirically determined for the Zeiss LSM 880 at 63x with Alexa Fluor 555 and may not transfer directly to different microscopes, fluorophores, or staining protocols. A more generalizable formulation would express the offset as a multiple of the background standard deviation, scaling automatically with the noise characteristics of each imaging system.

The *in vitro* sample size (n = 4-5 independent flow experiments per condition) was sufficient to detect differences in fragment area and junctional fragmentation ratio but may have been underpowered for total junctional area, where the overall ANOVA reached significance but no individual pairwise comparison survived Tukey’s correction. Larger sample sizes or additional experimental conditions would strengthen the statistical power of the pipeline’s output metrics.

### 4.2 *IN VIVO* PIPELINE LIMITATIONS

The ilastik pixel classifier was trained on five representative Z-stacks selected for high and low microglial content. While this training set was sufficient for the cohorts analyzed here, the classifier may not generalize to tissue processed with different staining protocols, imaged on different microscopes, or prepared with different fixation methods without retraining. The training process itself requires interactive annotation in the ilastik GUI, which took approximately one to two hours for the dataset used in this study.

Vessel mask selection retains a user verification step: the pipeline presents multiple mask variants, and the user selects the most accurate one. This introduces a subjective element, though it is confined to a binary choice between pre-computed options rather than a freehand segmentation. In practice, the choice between mask variants was rarely ambiguous for a given image, but inter-user agreement was not formally assessed.

The TBI cohort (n = 2) served as a positive control for BBB disruption rather than an experimental group. While TBI mice showed the expected marker changes, the sample size precluded statistical inference for this cohort. A fully powered TBI group would strengthen the validation of the pipeline’s sensitivity to acute injury.

With n = 4 per group, a sensitivity analysis indicates that the study could only reliably detect large differences between cohorts. For ZO-1, the pooled residual standard deviation from the two-way ANOVA was approximately 9.1 percentage points, which at this sample size and α = 0.05 provides 80% power to detect a difference of approximately 18 percentage points between groups. A difference smaller than this would not reach statistical significance even if it were real. The absence of significant aged-to-AD differences should therefore not be interpreted as evidence that no AD-specific effect exists, only that any such effect was not large enough to detect at this sample size. This limitation is particularly relevant to the eNOS finding in the PFC, where AD mice differed significantly from young controls, but aged mice did not differ from either group, leaving open whether eNOS upregulation in this region is truly AD-specific or represents the upper end of an age-related continuum that the study was underpowered to resolve.

All mice across all four cohorts were female. Female APPswe/PSEN1dE9 mice exhibit more severe amyloid pathology than males [47], meaning the vascular changes reported here may represent an upper bound for this model. Whether the same marker profiles hold in male mice is unknown. The APPswe/PSEN1dE9 model also has limited cerebral amyloid angiopathy, due to its Aβ42-dominant production profile [8], as discussed in Sections 1.1 and 1.4, and the finding that aged and AD mice were statistically indistinguishable may partly reflect this limited vascular amyloid burden rather than a true absence of AD-specific BBB changes.

All cohorts were analyzed at a single time point (3 months for young and TBI, 18 months for aged and AD). Multi-time-point cross-sectional comparisons would be needed to determine whether the vascular changes detected here are progressive.

Immunohistochemical quantification captures protein expression but not functional state. The increased eNOS coverage in AD mice should therefore be interpreted as evidence of altered endothelial phenotype rather than a direct measure of NO output. Increased protein expression may reflect compensatory upregulation of a dysfunctional (uncoupled) enzyme, with opposing consequences for barrier permeability depending on whether the enzyme produces protective NO or harmful reactive oxygen species [27]. Functional assays measuring NO output or enzyme coupling state would be needed to determine the biological significance of the eNOS coverage changes. The pipelines developed here quantify protein localization and coverage but do not report fluorescence intensity as an outcome variable. Intensity-based quantification would provide a complementary measure of protein expression level, but achieving reproducible intensity measurements across images requires tightly controlled confocal laser settings, antibody concentrations, and staining conditions that were not standardized across all imaging sessions in this study. Intensity quantification and complementary Western blot validation are underway but beyond the scope of this manuscript.

eNOS images were acquired at mixed magnifications and immersion media (20x with a 0.8 numerical aperture air objective and 40x with a 1.4 numerical aperture oil-immersion objective) across cohorts. Because the pipeline computes percent area coverage using per-image thresholding, the metric was normalized to each image’s own signal distribution, and coverage values did not differ systematically by imaging condition within cohorts where both magnifications were represented. ZO-1 and ICAM-1 were each acquired at a consistent magnification and immersion medium across all cohorts.

The use of female-only cohorts, a single AD model, and a single time point are all addressable by extending the pipelines to additional experimental conditions, as described in Section 4.3.

### 4.3 FUTURE DIRECTIONS

Automated junction detection via machine learning could eliminate the manual tracing step in the *in vitro* pipeline. The ilastik pixel classification framework already used for vessel-microglia separation in the *in vivo* pipeline could be trained to distinguish junctional ZO-1 from cytoplasmic ZO-1 based on local intensity patterns and spatial context. A classifier trained on the manually traced images from this study could serve as a starting point, with the existing manual tracings providing ground truth labels for training and validation.

Three-dimensional volumetric fragment analysis would address the limitation of collapsing Z-stacks into maximum intensity projections. By performing connected component analysis on the full Z-stack, fragments could be measured as volumes rather than areas, preserving axial continuity that the projection discards. To our knowledge, no existing tool performs 3D morphometric analysis of tight junction fragments, making this a novel direction for future development.

The modular design of the *in vivo* pipeline, where each marker is processed independently against a shared reference channel mask, means that additional markers can be incorporated without modifying the core analysis framework. Because the mask and the protein of interest are processed as independent channels, the same workflow applies to any fluorescent marker measured against any binary mask, not just the markers and lectin channel used here. Potential applications include quantifying pericyte coverage on vessels, astrocyte endfeet contact with the vasculature, or immune cell infiltration within defined tissue regions. The accuracy of the lectin-derived vessel mask could also be validated by co-staining a subset of sections with PECAM (CD31), which labels endothelium without microglial cross-reactivity, or with Iba1 to provide ground truth microglial labels for evaluating the ilastik classifier. The pipeline could also be applied to human postmortem brain tissue, where tissue autofluorescence and background staining are generally lower than in mouse tissue, potentially improving both the classification and thresholding steps.

Applying the *in vivo* pipeline to AD models with greater vascular amyloid burden would test whether AD-specific BBB changes emerge when cerebral amyloid angiopathy is more prominent. The Tg2576 model, which carries only the Swedish APP mutation, produces a higher ratio of Aβ40 to Aβ42 and develops more extensive vascular amyloid deposition. The 5xFAD model carries five familial AD mutations and exhibits aggressive plaque pathology with both parenchymal and vascular components. Either model may produce vascular marker changes that differ from the aging-dominated pattern observed in the APPswe/PSEN1dE9 mice and would more directly test whether AD-specific BBB dysfunction can be distinguished from normal aging at the protein level. Extending these analyses to male cohorts and to additional time points would further clarify the contributions of sex, age, and disease to the vascular marker profiles reported here. Future studies should also standardize imaging parameters, including magnification and immersion medium, across all markers during data collection. The eNOS coverage increase in the AD PFC raises the question of whether the additional protein is functional. Staining for phosphorylated eNOS (Ser1177) would distinguish between compensatory upregulation of functional enzyme and accumulation of uncoupled protein. Co-staining for VE-cadherin and eNOS in vivo would test whether eNOS upregulation corresponds to altered VE-cadherin localization at endothelial junctions, directly testing the uncoupling hypothesis discussed in Section 3.2.2.2. An in vitro BBB model could connect eNOS protein expression to nitric oxide output under controlled conditions. Administering exogenous Aβ *in vivo* or *in vitro* could also test whether direct amyloid exposure produces vascular marker changes detectable by either pipeline, independent of the transgenic background.

Similarly, the *in vitro* fragmentation pipeline could be applied to cells exposed to different stressors beyond GCX KD, such as inflammatory cytokines, shear stress, or pharmacological treatments, to determine whether different stressors produce distinct fragmentation phenotypes, which this pipeline would be able to quantify. The HBMECs analyzed in this study were cultured under physiological shear stress, and future work could examine whether static culture or pathological shear profiles produce distinct fragmentation signatures.

## 5. CONCLUSION

This study developed two image analysis pipelines for quantifying BBB-associated vascular markers *in vitro* and *in vivo*. Both pipelines are designed so that manual decisions are confined to upstream steps where investigator judgment is required, and all downstream measurement is automated. Once per-image inputs are defined (junction tracings for the *in vitro* pipeline; vessel mask selections, ROIs, and threshold adjustments for the *in vivo* pipeline), all downstream analysis is fully deterministic. Re-running the pipeline on the same inputs produces identical output, eliminating the inter-run variability inherent in manual scoring and categorical classification approaches. Both pipelines also reduce analysis time from minutes per image to seconds, with cumulative savings on the order of days across the datasets analyzed here.

The *in vitro* pipeline quantifies ZO-1 tight junction fragmentation through fragment-level morphometric analysis. Applied to HBMECs under GCX KD conditions, the pipeline detected significantly reduced fragment area and junctional fragmentation ratio in both CD44 KD and SDC1 KD cells relative to controls, providing continuous-variable measures of junction disruption that categorical scoring approaches cannot resolve. The fragment-based approach requires manual junction tracing but does not require automated cell detection or closed-perimeter identification, making it applicable to any preparation where junctions can be visually identified.

The *in vivo* pipeline quantifies vascular marker expression within lectin-defined vessel boundaries using ilastik-based pixel classification and FIJI macro-based colocalization analysis. ilastik addresses a problem that existing tools have not solved: discriminating two cell types labeled by the same fluorescent marker in a single channel. Prior machine learning approaches to microglial and vascular segmentation rely on dedicated channels where only one cell type is present. This approach operates entirely within the lectin channel, using morphological and textural features across planes to distinguish vessels from microglia without an additional stain. Applied across four mouse cohorts and three brain regions, the pipeline detected cohort- and region-specific differences in all three markers, with the findings detailed in Section 3.2. The marker-agnostic design of both pipelines, combined with their public availability on GitHub, supports their adoption by other laboratories for quantifying vascular markers beyond the specific systems analyzed in this study.

The two pipelines operate at different scales. The *in vitro* pipeline captures the spatial pattern of tight junction disruption at sub-cellular resolution, while the *in vivo* pipeline captures how that disruption is distributed across brain regions. Together, they show that semi-automated image analysis can extract reproducible, continuous-variable measurements from fluorescence microscopy data across both *in vitro* and *in vivo* BBB models.

## 7. STATEMENTS & DECLARATIONS

## Acknowledgements

We thank Prof. Abbas Yaseen and Chang Liu for providing the aged and AD mouse cohorts. We thank Eric Brengel of Prof. Craig Ferris’ laboratory for performing the TBI procedures and for providing the TBI cohort and a portion of the young cohort. We thank the laboratory of Prof. Chiara Bellini for providing the remainder of the young cohort. Finally, we thank Nicholas Mullikin for introducing us to ilastik pixel classification and providing initial instruction on its use, Lucas McCauley for assistance with tissue processing, and Prof. Abigail Koppes for valuable discussions and feedback that informed this work.

## Funding

This work was supported by Northeastern University start-up funds and the National Science Foundation CAREER Award [CMMI 1846962] to EEE. Support was provided by the National Science Foundation Graduate Research Fellowship (1938052) and the American Heart Association Predoctoral Fellowship (24PRE1188685) to NRO. Additionally, RLP was supported as a Sarafan ChEM–H Institute Scholar, MAC3 Impact Philanthropies Faculty Fellow, Knight Initiative Faculty Fellow, and Terman Faculty Fellow at Stanford University.

## Data Availability

The confocal image data underlying this report is available within the main text and publicly accessible in the Dryad repository at https://datadryad.org/. The Python codes used to implement image analysis pipelines are publicly available on GitHub. The *in vitro* pipeline is available at https://github.com/peckbe/junction-fragmentation-analysis, and the *in vivo* pipeline at https://github.com/peckbe/ilastik-fiji-colocalization. Both are published under the MIT license.

## Competing Interest Disclosures

The authors declare that there are no conflicts of interest to disclose.

## Author Contributions

- Research conception and design of experiments: BDP, NRO, CFF
- Data collection: BDP, NRO, CFF
- Processing, analysis, and interpretation of data: BDP
- Preparation of figures: BDP
- Drafting of the manuscript: BDP
- Editing and revising the manuscript: BDP, NRO, RLP, EEE
- Supervision and funding: CFF, RLP, EEE
- All authors approve of the final version of the manuscript.

## Notes

### Competing Interest Statement

The authors have declared no competing interest.

